# Sensing the shape of a surface by intracellular filaments

**DOI:** 10.1101/2024.11.18.624198

**Authors:** Handuo Shi, Jeffrey Nguyen, Zemer Gitai, Joshua Shaevitz, Benjamin P. Bratton, Ajay Gopinathan, Gregory Grason, Kerwyn Casey Huang

## Abstract

Understanding the mechanisms that dictate the localization of cytoskeletal filaments is crucial for elucidating cell shape regulation in prokaryotes. The actin homolog MreB plays a pivotal role in maintaining the shape of many rod-shaped bacteria such as *Escherichia coli* by directing cell-wall synthesis according to local curvature cues. However, the basis of MreB’s curvature-dependent localization has remained elusive. Here, we develop a biophysical model for the energetics of filament binding to a surface that integrates the complex interplay between filament twist and bending and the two-dimensional surface geometry. Our model predicts that the spatial localization of a filament like MreB with substantial intrinsic twist is governed by both the mean and Gaussian curvatures of the cell envelope, which strongly covary in rod-shaped cells. Using molecular dynamics simulations to estimate the mechanical properties of MreB filaments, we show that their thermodynamic preference for regions with lower mean and Gaussian curvatures matches experimental observations for physiologically relevant filament lengths of ∼50 nm. We find that the experimentally measured statistical curvature preference is maintained in the absence of filament motion and after a cycle of depolymerization, repolymerization, and membrane rebinding, indicating that equilibrium energetics can explain MreB localization. These findings provide critical insights into the physical principles underlying cytoskeletal filament localization, and suggest new design principles for synthetic shape sensing nanomaterials.

**Significance statement:** The protein MreB, a homolog of eukaryotic actin, regulates the shape of bacteria like *Escherichia coli* by guiding new cell-wall insertion based on local curvature cues. However, the mechanism by which a nanometer-scale MreB filament “senses” the micron-scale curvature of the cell wall has remained a mystery. We introduce a biophysical model of the energetics of twisted and bent filaments bound to curved surfaces, which predicts that localization of filaments like MreB is sensitive to both mean and Gaussian curvature. The model captures experimentally measured curvature enrichment patterns and explains how MreB naturally localizes to saddle-shaped regions without energy-consuming processes. Beyond cell shape regulation, our work suggests design principles for synthetic systems that can sense and respond to surface shape.

## Introduction

Understanding the mechanisms that enable sensing and control of cell shape in living organisms is a fundamental challenge. Known mechanisms are typically based on the regulation of proteins and their assemblies whose nanometer sizes are much smaller than the micron-scale dimensions of cells. A paradigmatic example is bacteria, whose cell shape is thought to be regulated by geometric control of the spatial pattern of membrane-bound cytoskeletal filaments (1). The energy landscape that describes the preferred localization of a stiff filament is a function of the mismatch between the geometry of the cell surface and the preferred filament conformation, which includes its propensity to curve and to twist (2). The local geometry of a surface is characterized by two principal curvatures, which can be expressed as a mean curvature (average magnitude) and a Gaussian curvature (the product of the two curvatures, whose sign indicates whether the surface is spherical or saddle-like). Intuitively, a protein or filament shaped like a one-dimensional strip with nonzero intrinsic curvature will prefer to bind to a surface along the direction that most closely matches that curvature.

Proteins whose structures are inherently two-dimensional can also localize in a manner determined primarily by the Gaussian curvature of the surface; for instance, amphipathic helices prefer Gaussian curvature (3) while *α*-synuclein generates negative Gaussian curvature (presumably due to its geometric preference) (4). Filament curvature and twist collectively interact with both aspects of curvature, yet many models of filament energetics focus on mean and Gaussian curvature separately without considering their covariation along the surface, which can be particularly high for common bacterial cell shapes such as rods.

Bacteria such as *Escherichia coli* maintain rod-like shape by actively remodeling their rigid peptidoglycan cell wall during growth (5). In many species, the pattern of cell-wall insertion is regulated by filaments of the actin homolog MreB (6-8). MreB forms short filaments that move along the cell periphery, directing where new cell wall material is incorporated, and its localization and movement are correlated with the local cell wall curvature (9-12). MreB is enriched in regions of both low mean and Gaussian curvature (9, 11), particularly in aberrantly shaped spheroplasts lacking a complete cell wall (12).

By mediating feedback between the local cell wall geometry and cell wall synthesis, MreB helps to robustly maintain a rod-like morphology at the cellular scale. Indeed, genetic depletion or chemical inhibition of MreB causes cells to lose their shape and eventually lyse. However, the precise nature of the coupling between cell geometry and MreB localization is still not fully understood (11); it is unclear whether MreB senses mean or Gaussian curvature directly, or if its localization is driven by its movement (13).

In many bacteria including *E. coli*, MreB forms an antiparallel double protofilament (14). While the double protofilament appears straight in crystal structures, molecular dynamics simulations have revealed the potential for it to adopt multiple twist states (15, 16). A previous biophysical model showed that the combined effects of the energy of membrane and filament bending, as well as a membrane pinning term that constrains the membrane to the shape of the cell wall, are sufficient to prescribe the orientation and length of a membrane-attached filament such as MreB (17). However, this model ignored variations in cell wall geometry away from a perfect cylinder and any contributions from filament twisting. Other models studying cell-shape homeostasis have also focused on filament bending (13, 18-20). Much like actin, the asymmetric cross-section and potential for twisting in MreB double protofilaments suggests that both bending and twisting should be considered on equivalent footing in energetic models. In actin, bending and twist are strongly coupled (21), and twisting plays an important role in the regulation of actin filament stability and assembly dynamics (22). A biophysical model considering the requirement that a twisted MreB filament straighten to align membrane binding sites with the surface successfully predicted changes in filament size and orientation across a range of MreB mutants with different intrinsic twists (16), suggesting that the anisotropy of surface binding and twisting are critical components of the energy landscape. However, this study again assumed that the cell surface was perfectly cylindrical; leaving unanswered the critical question of how variation in local surface geometry affects filament localization.

Here, we develop a model for filament localization based on the generic compatibility between the natural shape of filaments, their resistance to bending and twisting, and the arbitrary geometry of the surface to which they bind. We consider filaments across the range from pure bending to pure twisting and find that the coupling between twist and bending is a critical determinant of localization. Using molecular dynamics simulations to extract key mechanical parameters, our model quantitatively predicts an MreB enrichment profile consistent with experimental measurements for physiologically relevant filament lengths, and we demonstrate experimentally that enrichment is not dependent on MreB movement or a memory of previous binding patterns. Taken together, our findings indicate that the spatial localization of MreB is an equilibrium phenomenon that fundamentally depends on the covariance of mean and Gaussian curvature; that is, on the interdependence of curvatures within biologically relevant cell shapes and their fluctuations.

## Results

### Shape compatibility between twisted filaments and the surface to which they bind determines a curvature-dependent binding landscape

To determine how surface geometry and filament mechanics determine localization preferences, we first consider a generic model for the elastic energy of intrinsically twisted and bent anisotropic filaments that strongly adsorb to a surface of arbitrary mean (*H*) and Gaussian (*K*_*G*_) curvatures. As a simplified model of bacterial cytoskeletal filaments like MreB associating to am approximately cylindrical cytoplasmic membrane (whose rod-shaped geometry is determined by the cell wall), we consider short elastic rods with anisotropic local cross-section (Fig. 1A), whose wider dimension is assumed to mediate binding to the surface. We assume that the filament can bend easily in one direction (Fig. 1A) and possesses intrinsic curvature *k*_0_ in this direction, while it is intrinsically straight and substantially stiffer to bending in the orthogonal direction. The filament also possesses an intrinsic twist, *ω* _0_. The elastic energy of the filament can then be described by an anisotropic Kirchoff rod model:

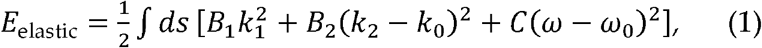

where *B*_1_ and *B*_2_ are the bending moduli in the two cross-sectional dimensions, with *B*_1_ and *B*_2_ corresponding to the bending in the direction parallel to the membrane binding surface and perpendicular to the membrane, respectively. *C* is the twist modulus. The filament’s uniform intrinsic curvature, *k*_0_ ≡ Ω_0_ sin *α*, in the wide dimension and intrinsic twist, *ω*_0_ ≡ Ω_0_ cos *α*, can be parameterized by a rotation per unit length Ω_0_ and angle *α*, where *α* parametrizes intrinsic geometries spanning from pure twist (*α =* 0) to pure bending (*α* = π/2) (Fig. 1A). The actual local shape of the filament is characterized by its curvatures *k*_1_ and *k*_2_ in the two cross-sectional dimensions and its twist *ω*, which are in general distorted away from the elastically preferred intrinsic values through interactions with the surface.

**Figure 1:**
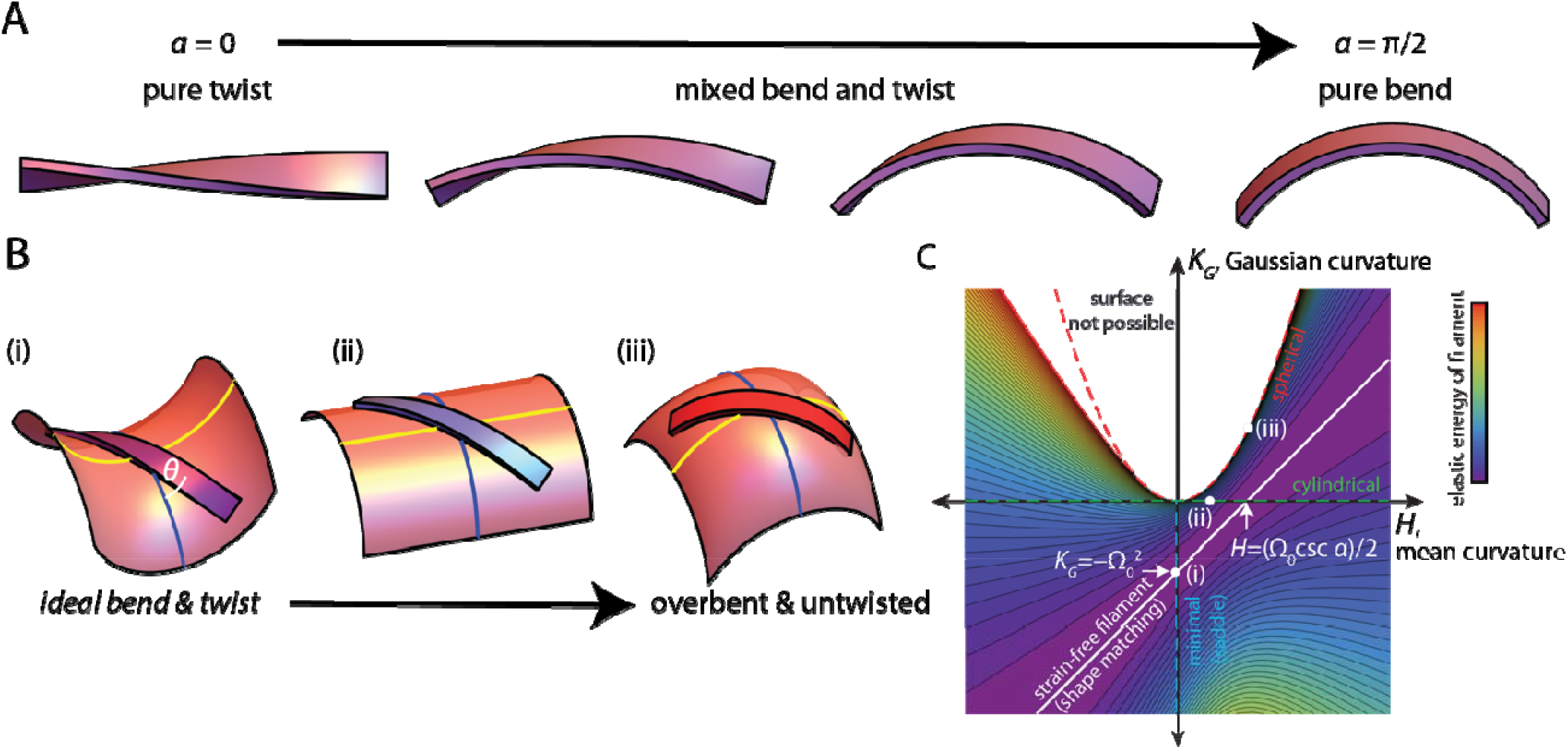
A general model for the localization of intrinsically twisted and bent filaments. A) Schematic of short filaments with intrinsic twist and bending. Parameterizes filament geometries spanning from pure twist () to pure bending (). For visualization purposes, the filaments are illustrated as flat bands with the wide cross-sectional dimension being the membrane-binding surface. B) For strong adsorption to curved surfaces shaped as a (i) saddle, (ii) cylinder, and (iii) sphere, filament shape is constrained by the local surface geometry. is the angle between the long axis of the filament and the principal surface direction with larger curvature. C) A representative energy landscape for a filament with 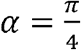. The white line represents a family of surface shapes with perfect geometric matching between the filament and surface for strain-free adsorption. White dots labelled (i), (ii), and (iii) correspond to the surface geometries in (B). Light blue region: geometries that are not possible; the white region directly to the left has very high elastic energy.

We consider a filament of fixed length *L* bound to a surface, such that *L* is larger than the transverse dimensions of the filament cross-section but substantially shorter than the local radii of curvature of the surface. The alignment of the wide face of the filament to the local surface normal constrains the filament (Supplemental Text; Fig. S1A) via

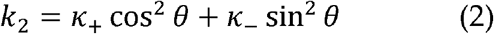

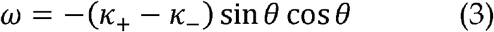

where *k*_±_ are the curvatures in the two principal directions of the surface at the point of filament contact, with *k*_+_ > κ_−_. Here, *θ* is the angle between the long axis of the filament and the principal direction with the larger curvature (Fig. 1B, left). We assume that *B*_1_ is sufficiently large that *k*_1_ = 0 (i.e., filaments follow geodesics), and that the filament is sufficiently short that variation in local curvature is negligible.

Using Eq. 1-3 and the parametrized forms of the intrinsic twist and curvature of the filament, the elastic cost of absorption per unit length *f* ≡ *E*_*elastic*_/*L* can be expressed as

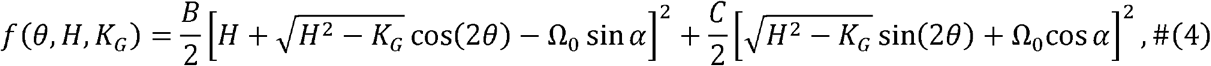

where *B* = *B*_2_ and *H* = (*κ*_+_ + *κ*_-_)/2 and *K*_*G*_ = *κ*_+_*κ*_*-*_ are the mean and Gaussian curvatures, respectively, at the location of surface contact. Upon binding, thermodynamics selects a filament orientation *θ* that minimizes the energetic cost to deviate from the preferred filament shape, leading to an elastic contribution that depends on local surface curvature defined by *f*_*\**_ (*H,K*_*G*_) = min_θ_ *f*(*θ,H, K*_*G*_). This relation defines an energy landscape of adsorption that varies with intrinsic filament geometry, parametrized by *α*. Fig. 1C shows an example of such a landscape for filaments with *α* = *π* /4, with nonzero intrinsic twist and bend. A critical and generic feature of these landscapes is the one-dimensional family of surface shapes for which *f* _*_ = 0 for all filament geometries (white line in Fig. 1C), indicating a perfect fit between filament and surface geometry that allows strain-free adsorption. The *f* _*_ = 0 condition can be determined directly from Eq. 4 (SI) to yield a linear relationship between Gaussian and mean curvature for surfaces that fit the filament perfectly:

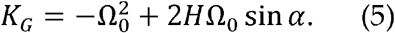

Combinations of mean and Gaussian curvature satisfying Eq. 5 correspond to geometries with surface-imposed curvature and twist that perfectly match the preferred bend and twist, respectively, of bound filaments. Any deviation of the surface curvatures from this perfect fit condition results in strain and increasing elastic energy of the bound filament. As an example, Fig. 1C highlights three locations in the curvature landscape corresponding to a filament with *α* = π/4 bound to a (i) saddle, (ii) cylinder, and (iii) sphere (Fig. 1B). For the saddle with zero mean curvature, there is a specific Gaussian curvature for which the elastic energy in Eq. 4 is zero (i.e.,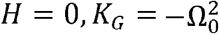), while the same filament bound to a cylinder and sphere experiences increasing elastic energy due to overbending and untwisting away from its preferred shape (Fig. 1B). Note that, independent of filament shape, perfect fit is generically possible for all values of mean curvature, yet the location of the perfect fit is skewed toward negative Gaussian curvature values (Fig. S1B), implying a generic preference for filament binding to surfaces with negative Gaussian curvature. For small *α* in particular, negative Gaussian curvature is favored due to the existence of straight (i.e., asymptotic) lines along which the surface normal twists. Thus, the shape of strongly binding filaments selects surface geometries with a specific linear relationship between mean and Gaussian curvature.

### Mean and Gaussian curvature are strongly correlated for rod-shaped bacterial cell geometries

To analyze localization along the membrane of rod-shaped bacteria such as *E. coli*, away from the approximately spherical poles, we consider an idealized model of local morphology: a bent cylindrical tube with a cross-sectional radius *r* whose centerline has a radius of curvature 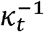 (i.e., a section of a toroid). The distribution of mean and Gaussian curvatures for this family of surfaces follows the simple linear relation

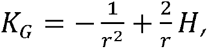

where for a given degree of bending, the range of variable curvature around the tube spans 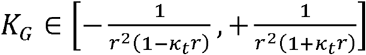. Hence, for the cylindrical body of a rod-shaped bacterium with fixed local radius but variable local curvature, the curvature distribution falls along the same line in curvature space and the range of Gaussian curvature values sampled increases in regions of the rod-like surface with high curvature (Fig. 2A).

**Figure 2:**
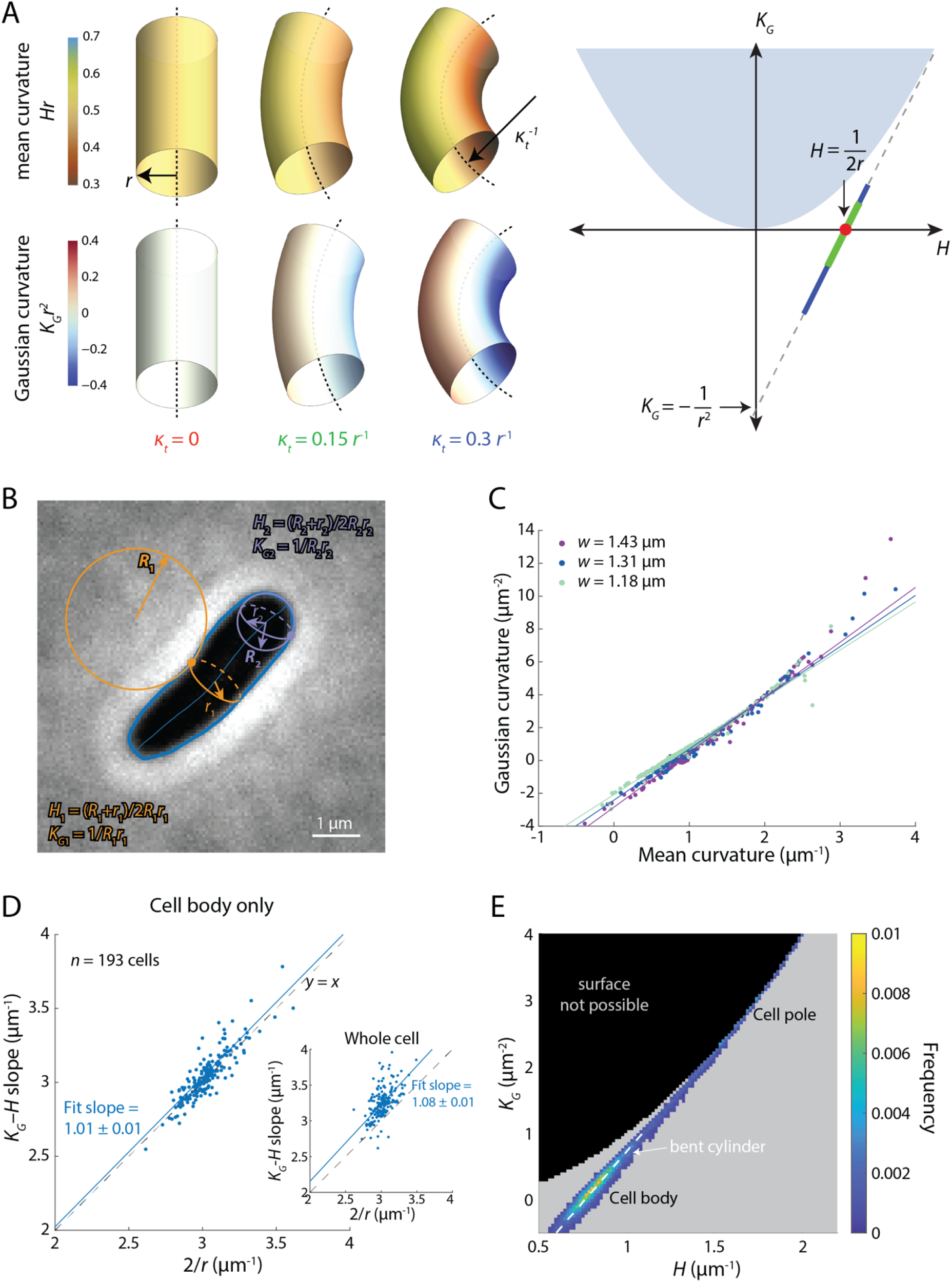
The cell body of a rod-shaped *E. coli* cells is approximately toroidal. A) Left: In an idealized model, the cell body of a rod-shaped cell can be approximated as a toroidal section with cross-sectional radius *r* and centerline radius of curvature 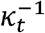. Right: For this family of surfaces, the accessible mean and Gaussian curvatures fall along a common line in curvature space regardless of *K*_*t*_, and toroids with larger *K*_*t*_ sample larger ranges of curvature values as shown by the red dot (*K*_*t*_ = 0), and green and blue (*K*_*t*_ = 0.15 *r* ^−1^ and *K*_*t*_ = 0.3 *r*^−1^, respectively) lines. Light blue region: geometries that are impossible. B) A representative phase-contrast image of an exponentially growing *E. coli* cell. The principal curvatures along the cell surfaces were estimated by assuming azimuthal symmetry (Methods). Blue: cell contour and midline. Purple and orange: estimated curvatures for two representative points along the cell contour. C) For several cell contours analyzed as in (B), mean and Gaussian curvatures were highly correlated, similar to the idealized bent tubes in (A). Exceptions were in regions with high curvatures corresponding to the hemispherical cell poles. As expected from (A), wider cells (with larger cross-sectional radius *r*) had larger slopes relating Gaussian and mean curvatures. D) Across a population of cells, the slope of the linear fit between Gaussian and mean curvatures measured along the cell body was approximately 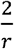, as expected from (A). Inset: this relationship was altered by inclusion of the cell poles, also as expected. E) The distribution of mean and Gaussian curvatures across a population of cells. Despite cell-to-cell variation, mean and Gaussian curvatures occupied a defined, highly constrained region, with most points falling around 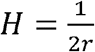 and *K*_*G*_ = 0, corresponding to the cylindrical cell body, and another subset of points with high *H* and *K*_*G*_ corresponding to the hemispherical poles. White dashed line: theoretical calculation based on a curved toroid as in (A). Gray represents geometries without data; black region is mean and Gaussian curvature combinations for which shapes are not possible. *n*=193 cells in (D) and (E).

To determine the extent to which actual rod-shaped bacterial cells are constrained to such toroidal geometries, we imaged a population of *E. coli* cells during log-phase growth and segmented the outline of each cell (Fig. 2B, Methods). Although it is difficult to precisely quantify bacterial geometry in three dimensions (23, 24), cells appear approximately rod-shaped; in particular, they do not exhibit regions along the cell body with large curvature compared to the inverse of the cylindrical radius. Thus, we assume that the cell centerline lies in the imaging plane (which is reasonable given the large mechanical pressure from the coverslip and the agarose pad during imaging), and estimate the principal curvatures of the cell surface based on the measured curvatures within and perpendicular to the imaging plane (Methods). For each point along the cell contour, we first obtained its osculating circle (with radius *R*) in the imaging plane. The radius of the osculating circle perpendicular to the imaging plane (*r*) was defined by the intersection of the local contour normal vector and the cell centerline (Fig. 2B). The corresponding mean and Gaussian curvatures of this point are 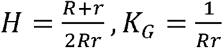. This strategy was previously determined to be reasonable for calculating the enrichment of MreB as a function of curvature (11). Similar to bent tubes (Fig. 2A), the relationship between *K*_*G*_ and *H* measurements in individual cells was approximately linear, except for regions with high *K*_*G*_ and *H* corresponding to the hemispherical poles (Fig. 2C). Indeed, across cells with different widths (hence different average cross-sectional radius *r*), we found that the slope of the line relating *K*_*G*_ and *H* was 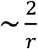 in the cell body (Fig. 2D). Aggregating across the entire population of cells, since cell width was narrowly distributed, most surface geometries had *K*_*G*_ = 0, representing cell bodies; a small subset near the poles had high *K*_*G*_ and *H* (Fig. 2E). Taken together, even with the presence of cell poles and local deformations away from rod-like shape, the mean and Gaussian curvatures in *E. coli* cells are strongly correlated and confined to a limited space that provides a framework for model predictions.

### Inferring the mechanical properties of MreB polymers from molecular dynamics simulations

We next sought to validate our model using the actin homolog MreB, which is essential in *Escherichia coli* (25). MreB forms short (diffraction-limited) filaments that bind to the inner surface of the cytoplasmic membrane and guide cell wall insertion (8). While it is challenging to experimentally measure the mechanical properties of MreB polymers, previous research has indicated that molecular dynamics (MD) simulations can provide quantitative estimates of mechanical parameters such as intrinsic bending and twisting and the associated moduli for filaments such as MreB (26, 27), the tubulin homolog FtsZ (28), and actin (29).

In crystal structures, MreB forms straight antiparallel double protofilaments (14). However, MD simulations revealed a broad range of potential conformational changes. MreB hydrolyses ATP (30), which influences the conformation of monomers and in turn is linked to the bending of a single protofilament (26). In a double protofilament, these nucleotide-dependent changes translate into a nucleotide-dependent twist (16). The bending and twist of MreB filaments are further affected by factors such as point mutations (16), binding to the MreB-interacting protein RodZ (27), and the presence of the small molecule MreB inhibitor A22 (16).

To capture the dynamics and mechanical properties of MreB over long time scales, we analyzed a 2.7-µs simulation of a 4×2 MreB double protofilament initialized from a crystal structure of *Caulobacter crescentus* MreB (PDB: 4CZJ, Fig. 3A, Table S1) (15). At each time step, we calculated the intrinsic curvature and twist of the polymer (Methods). Over extended intervals, we found that filament twist shifted among multiple steady states, corresponding to mean inter-subunit twist angles of 12.4°, 9.4°, and 4.3°, respectively (Fig. 3B), while filament bending was largely maintained at ∼2.4° in the direction perpendicular to the membrane-binding interface (Fig. 3C). *In vivo*, MreB filaments can be stable for tens of seconds (8), hence it is possible for them to experience the conformational changes observed *in silico*.

**Figure 3:**
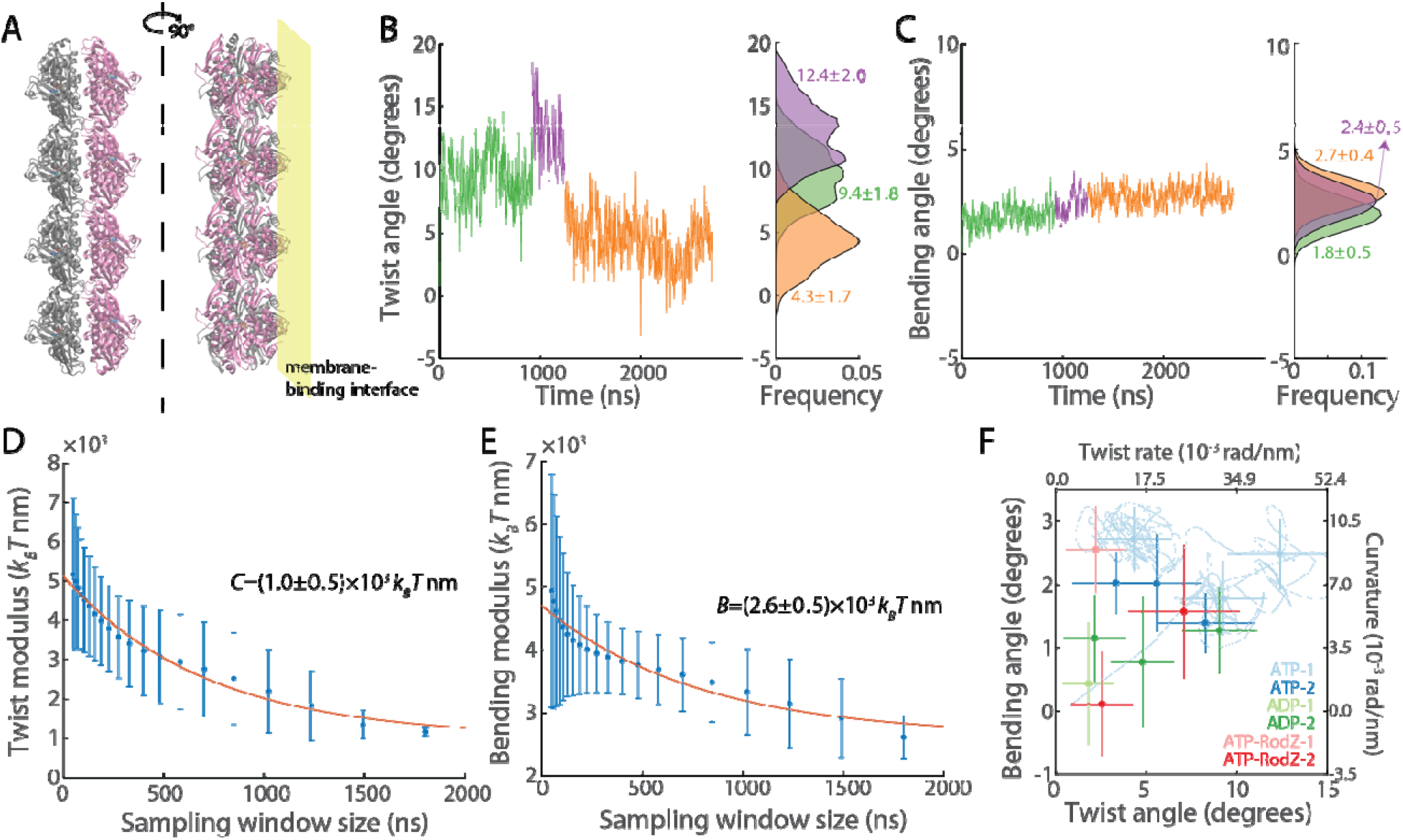
MD simulations reveal mechanical properties of MreB double protofilaments. A) Schematic of the initialization state for an MD simulation of a MreB double protofilament (based on PDB ID: 4CZJ). Yellow plane denotes where the membrane would be during adsorption but was not included in the simulations. B) In a representative simulation with MreB bound to ATP, the filament exhibited three distinct twist states. Left: raw trajectory colored by twist states. Right: histograms of the twist angles in each twist state. C) The bending angle from the simulation in (B). The three twist states exhibited similar bending angles. Left: raw trajectory colored by twist states in (B). Right: histograms of the bending angle in each twist state. D) The estimated twist modulus *C* monotonically decreased with larger sampling windows. We fit the estimates of *C* to a decaying exponential and used the asymptote to parameterize *C* in our biophysical model. E) The bending modulus *B* was similarly parameterized as in (D) using an exponential fit to estimated values of *B* across sampling windows. F) Across different simulation systems, MreB double protofilaments can adopt distinct twist states. Within each simulation, twist states were identified using a change-point analysis algorithm as in (B). Data are mean ± 1 standard deviation (S.D.). Light blue curve: raw trajectory of twist and bending angles for the simulation shown in (B) and (C).

We estimated the twist modulus of the MreB double protofilament from the MD simulation as 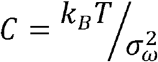, where *k*_*B*_ is the Boltzmann constant, *T* the absolute temperature, and *σ*_*ω*_ the standard deviation of fluctuations in the twisting angle _*ω*_; such a strategy was successfully used to estimate the twist modulus of actin filaments (29). We estimated *σ*_*ω*_ and therefore *C* based on the twist angle trajectory (Fig. 3B) using different sampling-window durations. Similar to the actin study (29), the twist angle of the MreB protofilament varied more in larger sampling-window durations, corresponding to smaller values of *C* (Fig. 3D). In the case of actin, the experimentally measured twisting modulus was close to the estimated value from the largest sampling window (29), likely because larger sampling windows captured more physiologically relevant dynamics. We therefore extrapolated the estimates of *C* by fitting to an exponential curve, and found the baseline of the fit to be (1.0 ± 0.5) × 10^3^ *k*_*B*_T nm (Fig. 3D); this value is very close to the value *C* estimated from the largest sampling window (1.1 × 10^3^ *k*_*B*_*T* nm). Using a similar approach, we estimated that the bending modulus of MreB is *B*= (2.6 ± 0.5) × 10^3^ *k*_*B*_*T* nm (Fig. 3E).

Since MreB protofilament conformation can be affected by factors like nucleotide binding and interacting proteins (16, 27), we also analyzed other simulations to capture the ranges of twist and bending angles for MreB double protofilaments bound to ADP or RodZ (Table S1). Since MreB double protofilaments can exhibit multiple twist states, we identified distinct twist states in each simulation using a change-point analysis algorithm (31, 32), and calculated twist and bending angles for each state (Methods). Across these simulations, MreB protofilaments exhibited inter-subunit twist angles ranging from 1.8 ° to 12.4° and bending angles from 0.1° to 2.7° (Fig. 3F); there were no obvious correlations between twist and bending (Fig. S2). Given that MreB monomer length in the filament direction is ∼5 nm (14), these values correspond to twist per unit length *ω* = 0.0063–0.0433 rad/nm, and curvature *k* = 0.0004–0.0095 rad/nm for MreB double protofilaments. These parameters provide the foundation for predicting MreB localization using our filament energetics model.

### Model prediction of MreB curvature enrichment

Armed with measurements of local geometries along *E. coli* cells (Fig. 2B) and estimates of the conformational and mechanical properties of MreB protofilaments (Fig. 3B-F), we used our filament mechanics model to predict the localization preference of MreB filaments in cells and compared these predictions to experimental observations of MreB localization. We acquired fluorescence images of MreB localization along the cell surface using an *E. coli* strain in which the only copy of *mreB* is a fusion to the fast-folding fluorescent protein msfGFP. Consistent with previous studies (9, 11, 27), we found that MreB is enriched at lower mean curvatures (Fig. 4A).

**Figure 4:**
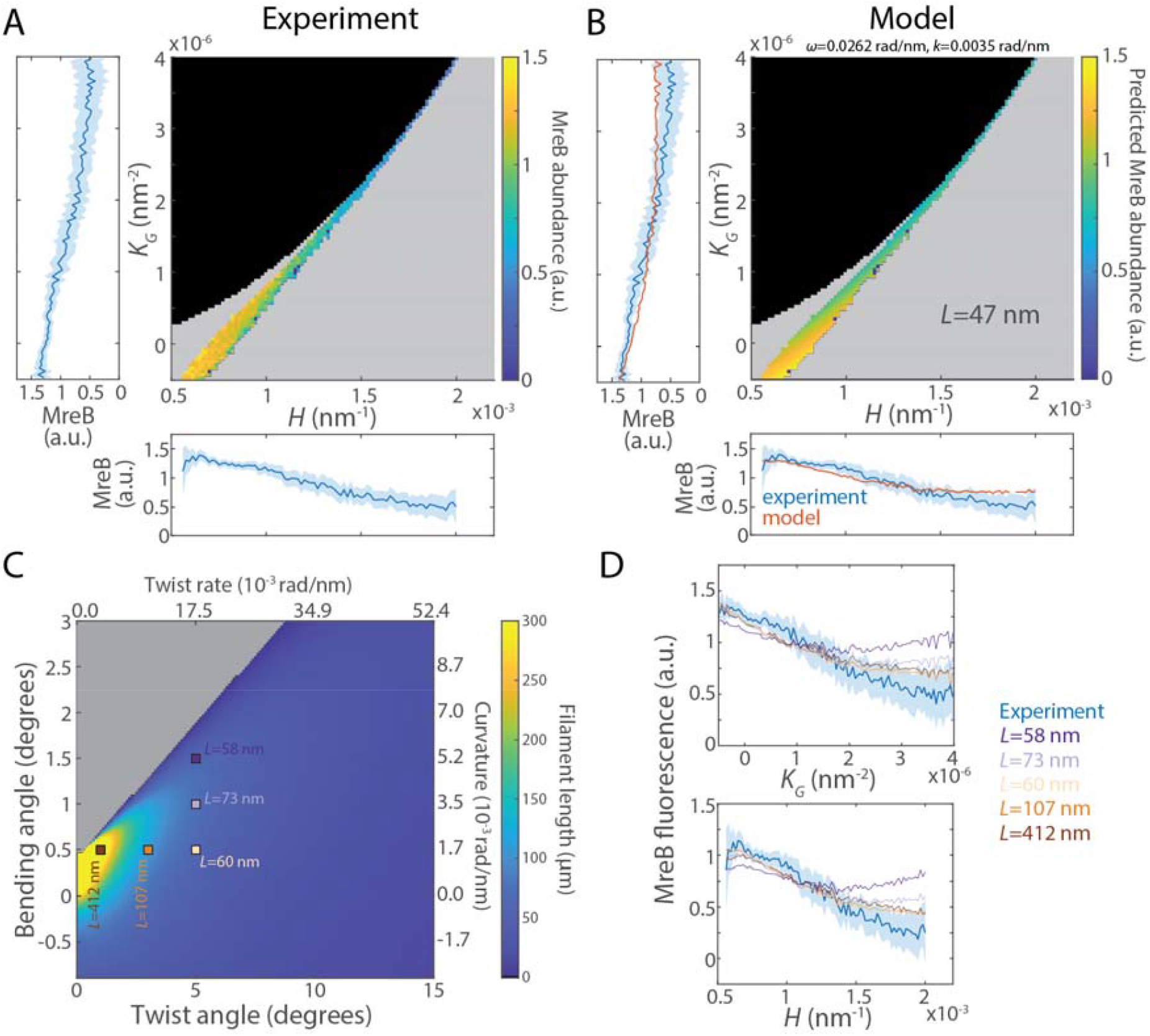
Model predicts biologically relevant MreB filament lengths. A) Experimental measurements of MreB enrichment binned by mean and Gaussian curvature in log-phase cells. Bottom histogram: MreB fluorescence distribution binned by mean curvature. Left histogram: MreB fluorescence binned by Gaussian curvature. Shaded regions denote 95% confidence intervals estimated by bootstrapping. Gray represents geometries with no data; black region is mean and Gaussian curvature combinations for which shapes are not possible. B) Fitting of experimental data in (A) to our model. The model recapitulates MreB enrichment at low curvatures and MreB depletion at high curvatures. The only free parameter is MreB filament length, which the best fit estimates at ∼47 nm. C) Phase diagram of predicted MreB filament length across bending and twist angles. Gray region: for these parameters, the model predicts that MreB should be enriched at regions of high rather than low curvature and thus the filament length for the best fit is zero. D) A subset of fits from (C) with different estimates of filament length. Although varying bending and twist angles changes the quantitative estimate of MreB filament length, the energy landscape remains qualitatively similar and the model still predicts preferential localization of MreB to lower mean curvatures, consistent with experiments.

To compare model results with experimental measurements, we selected intermediate values for twist and bending angles (twist angle 7.5° and bending angle 1.0°, corresponding to *ω* = 0.0262 rad/nm, *k* = 0.0035 rad/nm, Fig. 3F), and calculated the landscape of MreB elastic energy per unit length at each accessible combination of mean and Gaussian curvatures following Eq. 4. We assumed that MreB filaments always fully bind to the membrane, and therefore that membrane binding energy is constant. Thus, at each geometry defined by mean and Gaussian curvatures *H* and *K*_*G*_, we obtain the minimized local energy *E*_*_ (*H, K*_*G*_,*L*) = *L*min_*θ*_ *f*(*θ, H,K*_*G*_), where *L* is the MreB filament length, which we assumed for this analysis to be constant. Within cells, MreB localization preference is therefore set by a Boltzmann distribution

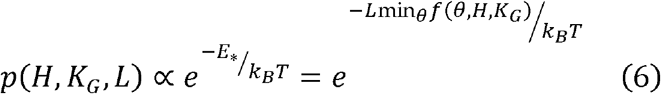

in which the only unknown parameter is *L*. We used Eq. 6 to predict the MreB localization preference as a function of curvature for a typical *E. coli* cell, and found a reasonable match to experimental measurements (Fig. 4B) with the filament length *L* = 47 nm. Importantly, the successful fit depends on the coupling of *H* and *K*_*G*_; if *K*_*G*_ is fixed at 0 (representing a perfect cylinder) then the model predicts that filament elastic energy would be higher at lower values of *H* (Fig. S3) and hence that MreB is enriched at higher mean curvature. Due to the geometric constraint imposed by the surface geometry, the model predicts that MreB filaments indeed deform from the strain-free conformation to bind to the membrane (Fig. S3). While our model predicts a general trend of MreB localization across the accessible range of mean and Gaussian curvatures that matches experiments, one quantitative difference is that at low curvatures, for a given mean curvature MreB is somewhat enriched at higher Gaussian curvature (Fig. 4A), opposite to the model prediction (Fig. 4B). We note that for fixed mean curvature, the regions with high or low Gaussian curvatures are accessible in cells across a range of widths (Fig. S4A). Wider cells typically have higher MreB concentrations (Fig. S4B), which potentially could affect MreB localization due to competition for binding sites but was not considered in our model.

We further explored how twist and bending angles affect the overall fit and estimated filament length *L* by performing the same fit with different *ω* and *k* values relevant for our MD simulations (Fig. 3F). For large *k* and small *ω*, representing the regime in which filament curvature dominates over twist, fits were particularly poor, as the elastic energy cost to unbend the filament becomes too large and hence higher cell surface curvatures are energetically more favorable (Fig. 4C, S4C). The preference for higher curvature holds generally in the extreme case of *ω* = 0, *k* > 0.0016 rad/nm (Fig. S4D,E), suggesting that intrinsic twist is critical for the experimentally observed MreB localization preference for lower curvatures. Nonetheless, for most combinations of *ω* and *k*, model fits were in reasonable agreement with experimental data, and predicted MreB filament lengths ranging from tens to a few hundred nm (Fig. 4C), largely consistent with MreB filament length measurements *in vivo* (16). For other values of the bending and twisting moduli *B* and *C*, while *L* varied quantitatively, the predicted curvature localization profiles remained similar (Fig. S5).

### Curvature-dependent localization of MreB is a statistical behavior independent of MreB motion

During steady-state growth, MreB filaments in *E. coli* (8) and *Bacillus subtilis* (6, 7) move in an approximately circumferential direction. The circumferential speed is correlated with the rate of cell-wall insertion and therefore bacterial growth. When cell-wall insertion in *E. coli* is disrupted by the PBP2 inhibitor mecillinam, MreB motion slows down in a dose-dependent fashion (8). It has been proposed that circumferential motion of MreB around a curved tube could by itself bias localization to negative curvature, since the inwardly curved side of the bent tube/toroid (with negative curvature) would collect a higher MreB density (19, 20) under the assumption that the curvatures on opposing sides of the cell body are anti-correlated, as in a perfectly curved toroid. We sought to test this critical assumption of anti-correlated curvature on opposing sides of the cell body. To minimize the confounding effects from cell poles and future division sites, we treated cells with the division inhibitor cephalexin. After excluding cell poles, we did not observe any correlations along the rod-shaped cell body (Fig. S6A), consistent with our previous observations (11). In fact, for ∼50% of the cell body, the curvatures on the opposing sides were either both positive or both negative, forming local bulges and indentations along the cell body (Fig. 5A). For cells not treated with cephalexin, future division sites generated some correlation when considering only negative contour curvatures, although again no correlation was observed across all curvatures (Fig. S6B). Therefore, although the curvatures on both sides of an idealized bent tube with a uniform cross-section are anticorrelated with each other (Fig. 2A), such a scenario does not apply in real cells, presumably due to fluctuations in the cell body geometry. Thus, circumferential motion alone likely does not explain MreB localization in *E. coli* cells.

**Figure 5:**
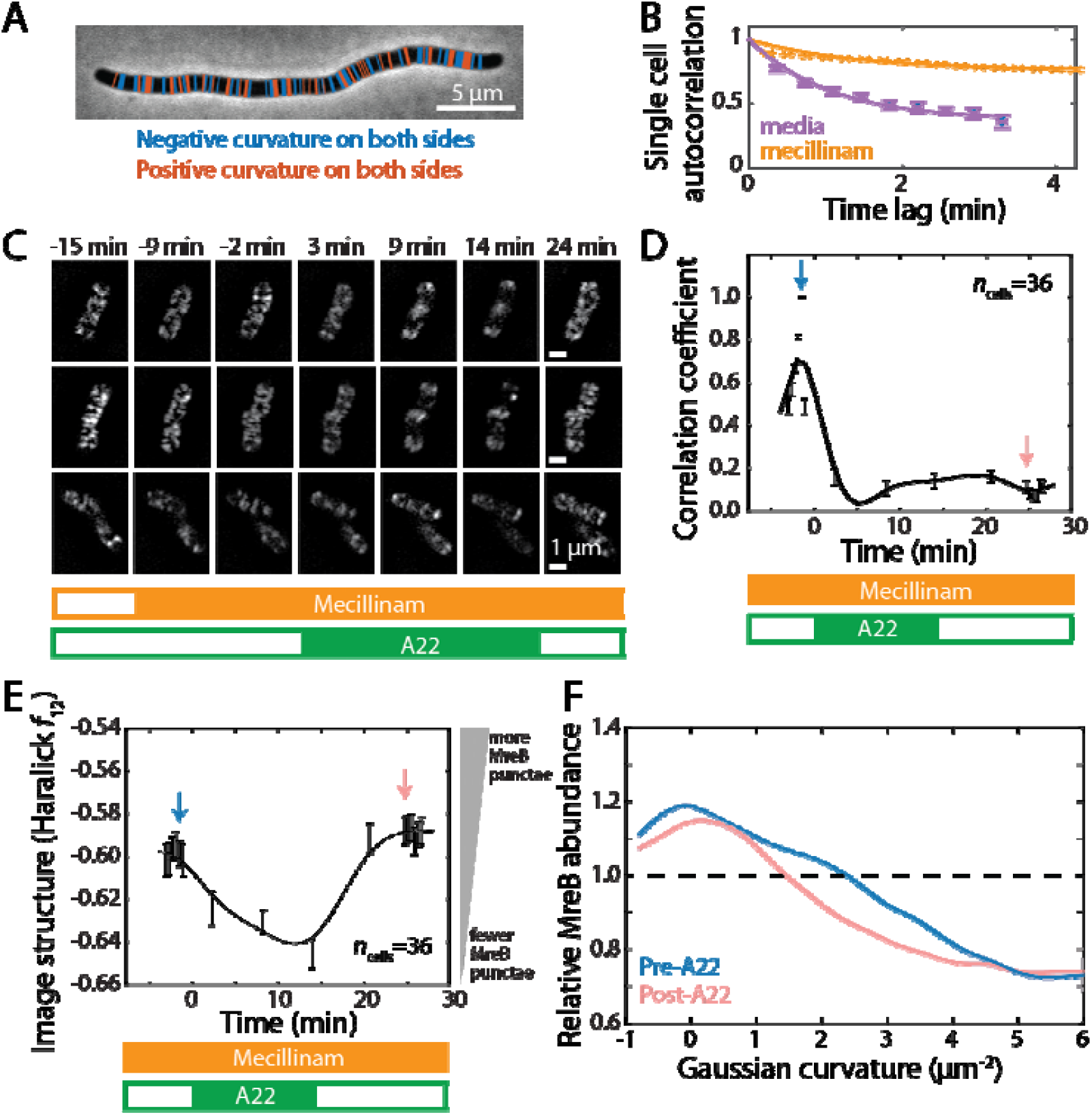
MreB enrichment is a statistical property of surface geometry, independent of its motion. A) In a filamentous *E. coli* cell treated with cephalexin, the in-plane curvatures along both sides of the cell body did not exhibit clear correlations, unlike an idealized bent toroid in which the curvatures on opposing sides always have opposite signs. Thus, local geometric variations in actual cells are shape deviations away from ideal toroids. B) When treated with mecillinam, MreB puncta largely halted movement, as indicated by the higher autocorrelation of single-cell fluorescence profiles. C) Representative mecillinam-treated cells that were transiently treated with A22. Bars at the bottom indicate the presence or absence of the two antibiotics in the microfluidic flow cell. *t* = 0 min corresponds to the time of A22 addition. D) Immediately following mecillinam administration, MreB motion halted, as indicated by the high correlation coefficient of intracellular fluorescence pattern relative to the reference pattern at *t* = -2 min. MreB then depolymerized when A22 was added, as indicated by the rapid decrease in the correlation coefficient. After A22 removal, the correlation coefficient remained low even though MreB repolymerized, indicating that the newly formed filaments bound to the membrane independent of previous localization sites. E) The status of MreB polymerization at each time point can be quantified using an image structure metric (Haralick *f*_12_). Higher values of Haralick *f*_12_ correspond to more structured images in which bright (dark) pixels are more likely to co-localize with other bright (dark) pixels (54). For MreB, higher values represent polymerization into filaments that appear as bright puncta. The dynamics indicate that addition of A22 resulted in rapid MreB depolymerization, and its removal led to MreB re-polymerization. F) Although MreB filaments localized to different regions of the cell before and after A22 treatment (D), the statistical distribution of MreB fluorescence as a function of Gaussian curvature remained similar. Blue and red arrows in (D) and (E) denote the two time points plotted here.

We therefore sought to test whether MreB exhibits a curvature preference in the absence of motion. We grew *E. coli* cells in a microfluidic flow cell until their growth rate reached a steady state, then treated the cells with mecillinam. Mecillinam effectively halted MreB movement, as quantified by a high autocorrelation of MreB fluorescence signal after treatment (Fig. 5B). While maintaining the presence of mecillinam, we treated the cells with A22 (33) to rapidly depolymerize the MreB filaments (Fig. 5C,D). After 15 min, we washed away A22, and MreB repolymerized onto the membrane; the re-formed MreB filaments did not move due to the continued presence of mecillinam (Fig. 5C,D). We then quantified the spatial distribution of MreB before and after A22 treatment. The precise locations of MreB filaments before and after A22 treatment were not correlated (Fig. 5E), indicating that MreB adopts a new pattern upon repolymerization. Nonetheless, the statistical profile of MreB enrichment as a function of curvature remained essentially the same (Fig. 5F), indicating that motion is not necessary for MreB’s bias to low mean curvature. Thus, we conclude that MreB localization can be explained by its energetic preference for local surface geometries with low mean and Gaussian curvature, which is dictated by the mechanical properties of MreB filaments.

## Discussion

Bacterial cells face a balance between maintenance of their cell size and shape and the benefits of modulating their morphology to adapt to changing growth conditions. Components that define the pattern of cell-wall synthesis must respond to any shape defects and reorganize downstream machinery to reinforce the preferred shape of a cell as it grows. Here, we demonstrated that the equilibrium energetics of MreB filaments are sufficient to recapitulate the observed localization of MreB to negative curvatures (Fig. 4), which acts as a negative feedback against bending fluctuations. Our results focused on exponentially growing cells, in which shape homeostasis may be particularly important; it will be intriguing to determine whether our energetic model is sufficient to predict MreB localization in other physiological states in which cell shape is altered.

Although the circumferential movement of MreB has been proposed to play a role in its observed localization to negative curvatures, we demonstrated that the statistical distribution of MreB localization is preserved even when motion is fully suppressed (Fig. 5), indicating that motion is not required and suggesting that MreB energetics alone may be sufficient for MreB localization. A mechanochemical model of cell wall growth (34) revealed that MreB movement can help to maintain a rod-like shape by smoothing out variations in cell-wall insertion when small morphological defects occur (8). However, it is challenging to rationalize the role of movement when cells severely deviate from a rod-like shape; for instance, in regions with variable cell width, movement can retain memory of the local geometry and hence could end up propagating such defects. A simulation framework that incorporates MreB movement and cell-wall dynamics under large perturbations and in abnormally shaped cells (12) will be informative in understanding the role of MreB movement in rod-shape regulation.

While our model can quantitatively predict most features of the observed pattern of MreB localization (Fig. 4), additional variables should be considered to comprehensively understand MreB properties. Our model consistently predicts higher MreB abundances in the polar regions than observed (Fig. 4B,D), possibly due to mechanisms actively excluding MreB from the poles, such as the enhanced concentration of anionic phospholipids (35). Moreover, the mechanical parameters we extracted from MD simulations (Fig. 3) may not completely reflect the properties of MreB. A recent study showed that physiologically relevant potassium concentrations substantially reduce MreB filament bending (36); MD simulations (including ours) typically do not incorporate potassium ions, so our estimates of bending and twist angles may be larger than *in vivo*, lowering estimates of filament length. Other MreB-interacting proteins like RodZ (24) and PBP2 (10) may also alter its energetic landscape.

Moving forward, more accurate biochemical measurements of MreB and protein structural insights provided by cryoelectron microscopy experiments and protein structure prediction tools like AlphaFold (37) will shed light on how MreB integrates with other cellular components to regulate cell shape.

Beyond the limitations of MD simulations, the statistical properties of MreB localization can realistically only be measured across many cells. MreB abundance can vary across cells (e.g., as a function of cell width; Fig. S4B), which can alter its localization and affect the accuracy of model predictions. To calculate the energy landscape of filament localization (Fig. 1), we assume that MreB filaments are of fixed length and short compared to variations in surface geometry, and that MreB strongly adsorbs to the membrane. In reality, variation in filament length due to (de-)polymerization would allow a single filament to sample different local geometries with modified energy landscapes. Moreover, weaker membrane adsorption could produce filaments that are only partially membrane bound, potentially with non-uniform morphologies in very long filaments (38), or could limit polymerization length due to energetic competition between polymerization and membrane binding (16). Previous models considering partial membrane binding have focused on cases with zero Gaussian curvatures (i.e., perfect cylinders) (16, 38). We expect that future enhancements to our model incorporating more realistic factors will further enhance predictions of cytoskeletal protein localization patterns, potentially in more complex geometries (12).

Is twist important beyond MreB? Evolutionarily, twist is a natural state of many polymers regardless of function (39, 40), and our study shows that a twisted polymer has intrinsic geometric preferences (Fig. 1, S1). Thus, cells may be able to appropriate polymers with suitable twist states for geometric sensing. In the case of MreB, point mutations can alter filament twist, which in turn changes cell morphologies and the pattern of MreB localization (9, 16). In addition, changes to filament bending and twist when bound to surfaces can exposure moieties responsible for other downstream processes (41), and filament untwisting could enhance parallel filament-filament interactions (42). These downstream effects could lead to autocatalytic amplification of the geometric sensing induced by filaments. The approach used in this study can also be generalized to other systems with intrinsic twist. For instance, twist can be introduced into filaments of the tubulin homolog protein FtsZ via point mutations, altering their localization and resulting in perturbations to cell division and shape (43). Future studies will provide insights on whether localization of other filaments is also determined by energetics or require factors such as movement (44, 45).

The general physical principles demonstrated in our work also empower *de novo* material designs. Our model of surface binding of filaments with intrinsic twist and curvature is generically applicable to engineering of shape sensors with novel functions. Engineered filaments with unique shapes, elasticities, and substrate binding properties can be used to measure and target specific material interfaces. Current protein design tools can rationally engineer such filaments (46), and their localization can be tested in complex geometries beyond bent toroids using microfluidic devices (47). Ultimately, twisted biopolymers such as DNA origami as well as shape-programmable polymer filaments and spiral nanosheet ribbons (48, 49) could be rationally designed to localize to distinct interfacial environments and even modify membrane geometry (50). Polymers and microfilaments that localize to mesoporous membranes may facilitate selective filtration, and proteins with geometric preferences can serve as scaffolds for tissue engineering. These applications highlight the opportunities for applying polymer physics models such as ours to geometry-sensing material functions.

## Acknowledgements

The authors thank the Huang lab for helpful discussions. The authors acknowledge support from NIH RM1 Award GM135102 (to K.C.H.), NSF Award EF-2125383 (to K.C.H.), a James S. McDonnell Postdoctoral Fellowship (to H.S.), NSF Awards DMR-2028885 and 2349818 (to G.M.G), NSF Awards HRD-1547848 and 2112675 (to A.G.) and partial support from CMMI-154857 (to A.G.). K.C.H. is a Chan Zuckerberg Biohub Investigator. This work was also supported in part by the National Science Foundation under Grant PHYS-1607611 and the hospitality of the Aspen Center for Physics.

## Methods

### Strains and media

Strains used in this study are *E. coli* MG1655 (CGSC #6300) and MG1655 *csrD::Km, mreB’::msfGFP-mreB’’* (11, 23). For experiments in Fig. 2B-E, 4A, and 5A, cells were cultured in lysogeny broth (LB; 10 g/L tryptone, 5 g/L NaCl, and 5 g/L yeast extract) and imaged at 37 °C on 1% agarose pads with LB. For experiments in Fig. 5B-F, cells were cultured at 37 °C in M63 minimal media [2 g/L (NH_4_)_2_SO_4_, 13.6 g/L KH_2_PO_4_, 0.0005 g/L FeSO_4_, and 0.25 g/L MgSO_4_·7H_2_O] supplemented with glucose and casamino acids, and imaging was performed in microfluidic channels constructed from two pieces of double-stick tape sandwiched between a coverslip and a glass slide. Cells were adhered to the surface by pretreating the coverslip with 0.1% poly-L-lysine solution for 5 min before adding cell suspension.

### Imaging

For experiments in Fig. 2B-E, 4A, and 5A, cells from exponentially growing cultures were placed on agarose pads and immediately imaged with a Nikon Ti-E inverted microscope (Nikon Instruments) using a 100X (1.40 NA) oil immersion objective and a Neo 5.5 sCMOS camera (Andor). Images were acquired using µManager v. 1.4 (51).

For experiments in Fig. 5B-E, images were acquired on a Nikon N-SIM microscope with a 100X (1.49 NA) objective and an Andor DU-897 EMCCD in Slide 3D-SIM reconstruction mode. For experiments in Fig. 5F, images were acquired on a custom monolithic aluminum microscope with a 100X (1.49 NA) objective (Nikon), an Andor DU-897 EMCCD, and custom LabView (National Instruments) software. Three-dimensional images were acquired with a *z*-stack step of 100 nm, and three dimensional shape reconstructions were performed using in house MATLAB software (https://github.com/PrincetonUniversity/shae-cellshape-public) (52).

### Single-cell segmentation and MreB enrichment analyses

For experiments in Fig. 2B-E, 4A, and 5A, cell contours were extracted from phase-contrast images using *Morphometrics* (53). The local curvatures along cell contours were calculated assuming azimuthal symmetry along the cell and that the cell centerline lies in the imaging plane. Under this assumption, the first principal curvature (*κ*_1_) is approximately within the imaging plane and was calculated by fitting an osculating circle to three neighboring contour points, then *κ*_1_ was smoothed along the cell contour using a moving Gaussian average. *κ*_1_ is negative if the osculating circle lies outside the cell contour, and positive otherwise. The second principal curvature (*κ*_2_) was estimated from the radius defined by the intersection of the local contour normal vector and the cell centerline (53); *κ*_2_ is always positive. Mean and Gaussian curvatures were calculated as 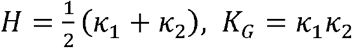. MreB intensities were measured along the cell contour as done previously (11) and normalized within each cell.

MreB enrichment profiles as a function of Gaussian curvature in Fig. 5F were calculated from three-dimensional measurements using code available at https://github.com/PrincetonUniversity/shae-cellshape-public. In brief, the shape of individual cells was reconstructed by minimizing the image difference between an observed focal stack and the forward convolution of a shape with the observed microscope blur function. At each iteration of the algorithm, individual surface elements (positions of vertices and their connectivity) are moved to reduce this image difference. Following these reconstructions, the relative MreB abundance was calculated as the weighted sum of MreB fluorescence as a function of surface curvature normalized by the surface area with that curvature. Values above 1 imply enrichment, or an average MreB concentration that is larger than would be expected from a uniformly distributed protein of the same intensity (52).

### Filament twist and bending

For each MD simulation frame, a local coordinate system was defined by three unit vectors (*d*_1_, *d*_2_, *d*_3_) as previously described (16). For an MreB double protofilament, *d*_3_ is largely parallel to the filament, *d*_2_ is perpendicular to the surface of MreB membrane binding, and *d*_1_ = *d*_3_ × *d*_2_. Twist angle is the rotation angle around *d*_3_. Bending is the rotational angle around *d*_2_. The bending angle around *d*_1_ was relatively small and fluctuated around zero.

### Identifying simulation twist states

Twist states were defined using the change-point analysis algorithm *Steppi* (31, 32). This algorithm identifies transitions between discrete states by assuming features of the noise in each state. For our analyses, parameters were chosen as follows (15): state level *µ* = unfixed, level slope *α* = 0, nearest-neighbor coupling *ϵ* = 0.6, noise “stiffness” *κ* = 0.2.

