## Supplemental Text for "Sensing the shape of a surface by intracellular filaments"

**Model of binding of a twisted filament to a surface relates energy to mean and Gaussian curvature**

We model filaments as anisotropic Kirchoff rods that have inextensible centerlines and cross-sections that remain planar and normal to their centerlines. They are described by a central curve $\mathbf{r}\left( s \right)$ and a local frame $\{{\hat{\mathbf{e}}}_{\boldsymbol{1}}\left( s \right),{\hat{\mathbf{e}}}_{\boldsymbol{2}}\left( s \right),\hat{\mathbf{t}}\left( s \right)\}$. Here, $\hat{\mathbf{t}}\left( s \right)=\partial_{s}\mathbf{r}$ is the tangent vector, and ${\hat{\mathbf{e}}}_{\boldsymbol{1}}\left( s \right)$ and ${\hat{\mathbf{e}}}_{\boldsymbol{2}}\left( s \right)$ define the material frame of the filament cross-section (Fig. S1). While the filament is represented schematically as a flat band with the wide cross-sectional dimension being the membrane-binding interface for ease of visualization, the actual cross-sectional dimensions do not affect the conclusions of our model. For concreteness, we define the ${\hat{\mathbf{e}}}_{\boldsymbol{1}}$ and ${\hat{\mathbf{e}}}_{\boldsymbol{2}}$ directions as aligned with the two dimensions of a ribbon-like local cross-section. This anisotropy allows us to capture two physical ingredients. First, we assume that the intrinsic shape is straight and perpendicular to the wide dimension (i.e., the preferred shape has zero intrinsic curvature) and the bending stiffness in this direction is significantly larger than in the orthogonal direction. Second, the cross-sectional anisotropy couples to the local (adhesive) interactions between the filament and the surface to which it is bound, such that interactions favor the alignment of a particular direction in the material frame with surface normal $\boldsymbol{N}$. Here, we assume that direction is ${\hat{\mathbf{e}}}_{\boldsymbol{2}}$, consistent with favorable contact between the surface and the wider direction of the cross section.

The local shape of the filament is dictated by its twist $\omega$ and curvatures $k_{1}$ and $k_{2}$ in the material directions:

$$\begin{aligned} \omega={\hat{\mathbf{e}}}_{\boldsymbol{2}}\cdot\left( \partial_{s}{\hat{\mathbf{e}}}_{\boldsymbol{1}} \right); k_{i}={\hat{\mathbf{e}}}_{\boldsymbol{i}}\cdot\left( \partial_{s}\hat{\mathbf{t}} \right)\#(1) \end{aligned}$$

where $i=1,2$. The elastic energy of the filament is

$$\begin{aligned} E_{\text{elastic}}=\frac{1}{2}\int ds\left[ B_{1}k_{1}^{2}+B_{2}\left( k_{2}-k_{0} \right)^{2}+C\left( \omega-\omega_{0} \right)^{2} \right], \#(2) \end{aligned}$$

where $B_{1,2}$ are the bending moduli in the two cross-sectional dimensions and $C$ is the twist modulus. We assume that the filament has uniform intrinsic curvature $k_{0}=\Omega_{0}\sin\alpha$ and intrinsic twist $\omega_{0}=\Omega_{0}\cos\alpha$, parameterized by rotation per unit length $\Omega_{0}$ and angle $\alpha$.

For a filament of fixed length $L$ bound to a surface such that $L$ is much shorter than the radii of curvature of the surface but much larger than the transverse dimensions of the filament cross-section, we parameterize the filament shape by expressing its local frame, $\{{\hat{\mathbf{e}}}_{\boldsymbol{1}}\left( s \right),{\hat{\mathbf{e}}}_{\boldsymbol{2}}\left( s \right),\hat{\mathbf{t}}\left( s \right)\}$, in terms of a local coordinate basis of the surface $\{{\hat{\mathbf{x}}}_{\boldsymbol{+}},{\hat{\mathbf{x}}}_{\boldsymbol{-}},\hat{\boldsymbol{N}}\}$, where ${\hat{\mathbf{x}}}_{\boldsymbol{\pm}}$ are the principal directions of the surface (Fig. S2) corresponding to principal curvatures $\kappa_{\pm}$:

$$\begin{aligned} {\hat{\mathbf{e}}}_{\boldsymbol{1}}=-\sin\theta{\hat{\mathbf{x}}}_{\boldsymbol{+}}+\cos\theta{\hat{\mathbf{x}}}_{\boldsymbol{-}}; {\hat{\mathbf{e}}}_{\boldsymbol{2}}=\hat{\boldsymbol{N}}; \hat{\boldsymbol{t}}=\cos\theta{\hat{\mathbf{x}}}_{\boldsymbol{+}}+\sin\theta{\hat{\mathbf{x}}}_{\boldsymbol{-}}, \boldsymbol{\#(}3) \end{aligned}$$

where the axes are defined such that $\kappa_{+}>\kappa_{-}$. We note that the alignment between the surface normal and the normal to the wide dimension of the filament is enforced (${\hat{\mathbf{e}}}_{\boldsymbol{2}}=\hat{\boldsymbol{N}}$) and that the angle between the filament tangent, $\hat{\boldsymbol{t}}$, and the larger principal curvature direction, ${\hat{\mathbf{x}}}_{\boldsymbol{+}}$**,** is taken to be $\theta$. This formulation enables us to express the principal curvatures in terms of the gradients of the principal direction unit vectors along the filament as

$$\begin{aligned} \kappa_{+}\cos\theta=\hat{\boldsymbol{N}}\cdot\left( \partial_{s}{\hat{\mathbf{x}}}_{\boldsymbol{+}} \right); \kappa_{-}\sin\theta=\hat{\boldsymbol{N}}\cdot\left( \partial_{s}{\hat{\mathbf{x}}}_{\boldsymbol{-}} \right). \#\left( 4 \right) \end{aligned}$$

Utilizing the expressions from Eq. 3 in Eq. 1, along with Eq. 4, and noting that ${\hat{\mathbf{x}}}_{\boldsymbol{-}}\cdot\left( \partial_{s}{\hat{\mathbf{x}}}_{\boldsymbol{+}} \right)={\boldsymbol{-}\hat{\mathbf{x}}}_{\boldsymbol{+}}\cdot\left( \partial_{s}{\hat{\mathbf{x}}}_{\boldsymbol{-}} \right)$, the filament twist and curvatures are related to the local surface geometry at $\boldsymbol{r}\left( s \right)$:

$k_{1}=\partial_{s}\theta+{\hat{\mathbf{x}}}_{\boldsymbol{-}}\cdot\left( \partial_{s}{\hat{\mathbf{x}}}_{\boldsymbol{+}} \right);k_{2}=\kappa_{+}\cos^{2} \theta+\kappa_{-}\sin^{2} \theta$;

$$\begin{aligned} \omega=-\left( \kappa_{+}-\kappa_{-} \right)\sin\theta\cos\theta. \#\left( 5 \right) \end{aligned}$$

We assume that $B_{1}$ is sufficiently large that $k_{1}=0$, but filaments are sufficiently short compared to the rotation rate of the principal curvature directions that any energetic costs of non-constant $\theta$ can be ignored.

The elastic cost of absorption per unit length $f\equiv E_{\text{elastic}}/L$ is then

$$\begin{aligned} f\left( \theta,H,K_{G} \right)=\frac{B}{2}\left[ H+\sqrt{H^{2}-K_{G}}\cos\left( 2\theta\right)-\Omega_{0}\sin\alpha\right]^{2}+\frac{C}{2}\left[ \sqrt{H^{2}-K_{G}}\sin\left( 2\theta\right)+\Omega_{0}\cos\alpha\right]^{2},\#\left( 6 \right) \end{aligned}$$

where ${B=B}_{2}$ and $H=\left( \kappa_{+}+\kappa_{-} \right)/2$ and $K_{G}=\kappa_{+}\kappa_{-}$ are the mean and Gaussian curvatures, respectively, at the location of contact with the surface. The elastic contribution to surface binding is obtained from the minimum of $f$ with respect to $\theta$ for fixed local surface shape,

$$\begin{aligned} f_{*}\left( H,K_{G} \right)=\text{min}_{\theta}f\left( \theta,H,K_{G} \right). \#\left( 7 \right) \end{aligned}$$

There exists a one-dimensional family of surface shapes for which $f=0$, indicating perfect fit between filament and surface geometry. The $f_{*}=0$ condition can be determined by combining the conditions for zero bending, $\cos\left( 2\theta\right)=\left[ \Omega_{0}\sin\alpha-H \right]/\sqrt{H^{2}-K_{G}}$, and for zero twist, $\sin\left( 2\theta\right)=-\Omega_{0}\cos\alpha/\sqrt{H^{2}-K_{G}}$, to yield a linear relationship between Gaussian and mean curvature, which is notably independent of bend and twist stiffnesses:

$$\begin{aligned} K_{G}=-\Omega_{0}^{2}+2H\Omega_{0}\sin\alpha, (\mathrm{for} f_{*}=0). \#\left( 8 \right) \end{aligned}$$

To determine the energetics away from the perfect conformal fitting, we consider the torque equilibrium

$$\begin{aligned} \frac{\partial f}{\partial\theta}=-2B\sqrt{H^{2}-K_{G}}\sin\left( 2\theta\right)\left[ H-\Omega_{0}\sin\alpha\right]+2C\sqrt{H^{2}-K_{G}}\cos\left( 2\theta\right)\Omega_{0}\cos\alpha\\ +\left( B-C \right)\left[ H^{2}-K_{G} \right]\sin\left( 4\theta\right)=0. \#\left( 9 \right) \end{aligned}$$

For the special case of equal twist and bend stiffness ($B=C$), the equilibrium angle $\theta_{*}$ satisfies

$$\begin{aligned} \tan\left( 2\theta_{*} \right)=\frac{\cos\alpha}{H\Omega_{0}^{-1}-\sin\alpha}, \#\left( 10 \right) \end{aligned}$$

which is independent of Gaussian curvature. This solution gives the dependence of the elastic energy of adsorption on $H$ and $K_{G}$:

$$\begin{aligned} f_{*}\left( B=C \right)=\frac{B}{2}\left[ \sqrt{H^{2}-2H\Omega_{0}\sin\alpha+\Omega_{0}^{2}}-\sqrt{H^{2}-K_{G}} \right]^{2}. \#\left( 11 \right) \end{aligned}$$

Note that Eq. 11 gives the energetics of misfit for arbitrary surface curvature values (for the special case of $B=C$), and hence it also satisfies $f_{*}=0$ for the “perfect fit” condition in Eq. 8.
**Supplemental Tables**

**Table S1: MD simulation systems analyzed in this study.** All simulations were originally reported in Ref. (1).

| **Name** | **Ligand** | **Simulation time (µs)** | **Total twist states** | **Time intervals defining each state** |
| --- | --- | --- | --- | --- |
| ATP-1 | ATP, Mg^2+^ | 2.71 | 3 | State 1: 0–0.93 µs  State 2: 0.93–1.28 µs  State 3: 1.28–2.71 µs |
| ATP-2 | ATP, Mg^2+^ | 2.01 | 3 | State 1: 0–0.46 µs, 0.58–1.06 µs, 1.63–2.01 µs  State 2: 0.46–0.58 µs, 1.20–1.63 µs  State 3: 1.06–1.20 µs |
| ADP-1 | ADP, Mg^2+^ | 1.00 | 1 | State 1: 0–1.00 µs |
| ADP-2 | ADP, Mg^2+^ | 1.01 | 3 | State 1: 0–0.37 µs  State 2: 0.37–0.51 µs  State 3: 0.51–1.01 µs |
| ATP-RodZ-1 | ATP, Mg^2+^, RodZ | 1.01 | 1 | State 1: 0–1.01 µs |
| ATP-RodZ-2 | ATP, Mg^2+^, RodZ | 1.01 | 2 | State 1: 0–0.46 µs  State 2: 0.46–1.01 µs |

**Supplemental Figures**

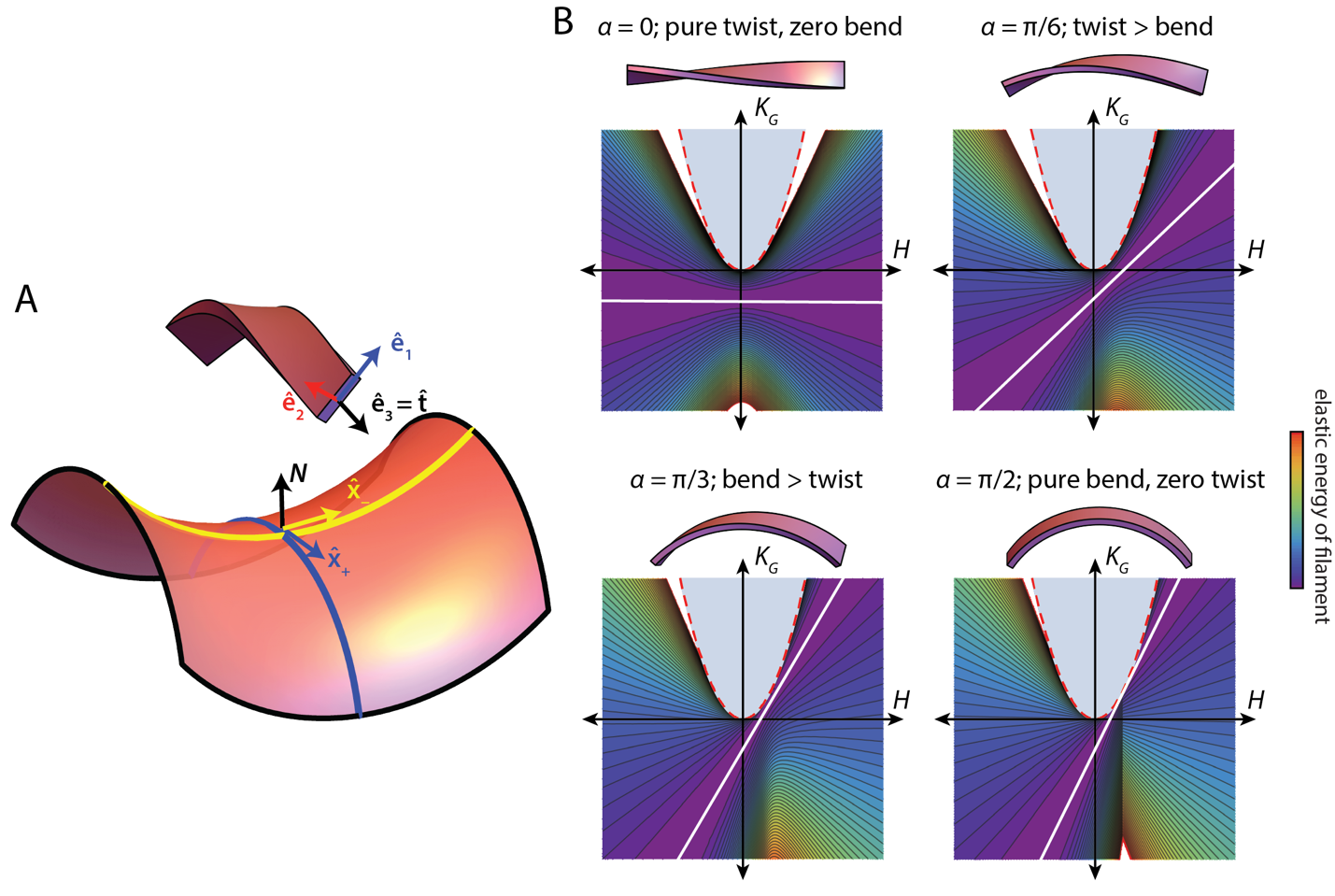

**Figure S1: Energetic landscapes for intrinsically twisted and bent filaments.**

A) Schematic of a twisted and bent filament binding to a curved surface.
B) The energy landscape for filaments with various values of $\alpha$. White lines represent a family of surface shapes that enable a perfect fit between the filament and surface for strain-free adsorption. Light blue regions: geometries that are impossible; adjacent white regions represent very high elastic energies.

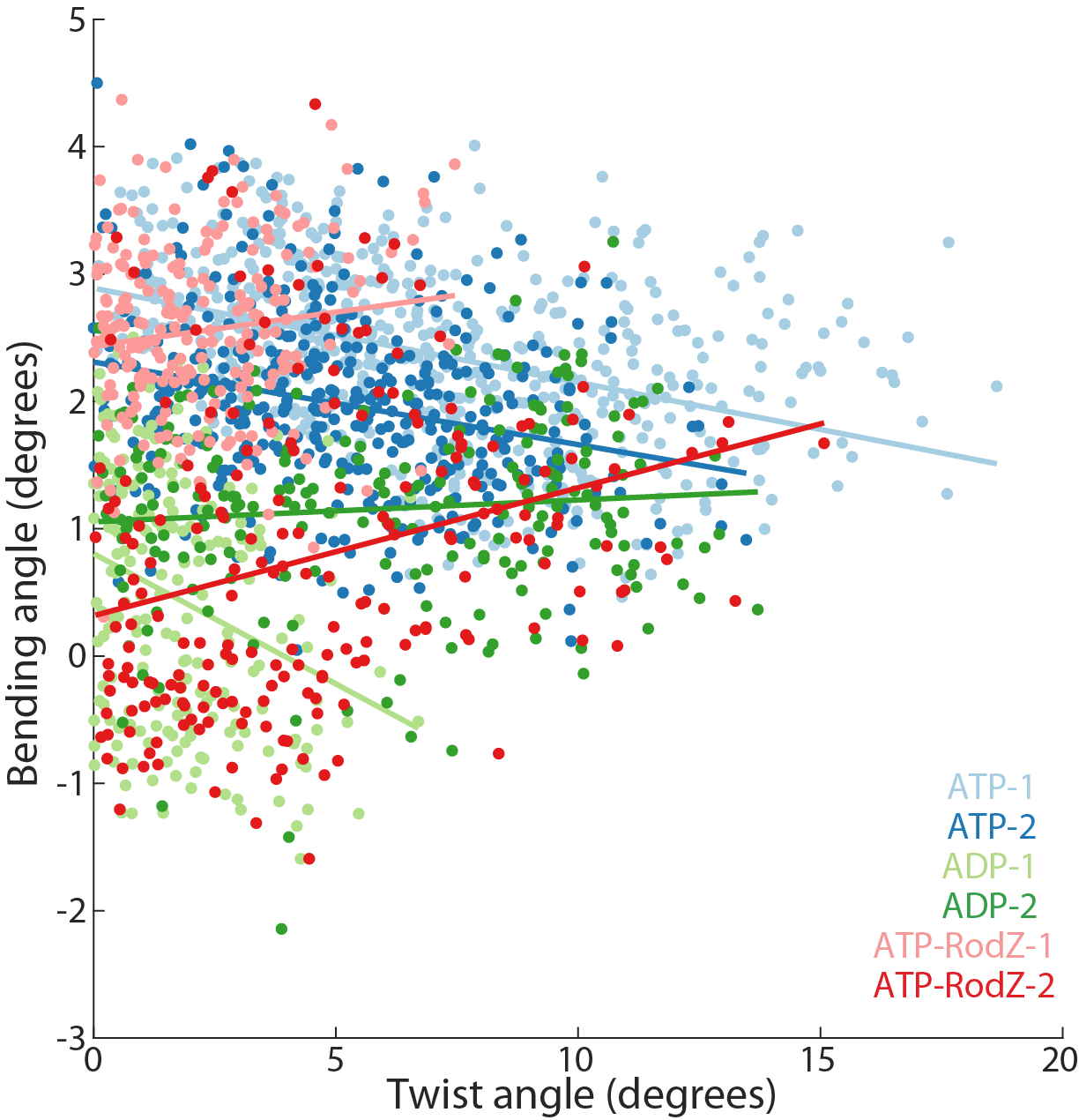

**Figure S2: MreB twist and bending angles are not obviously linked in MD simulations.** Scatter plot of twist and bending angles across all simulations analyzed in Fig. 3F. No systematic trends were observed between the two angles. Circles are raw data points from the simulations, and lines are best linear fits.

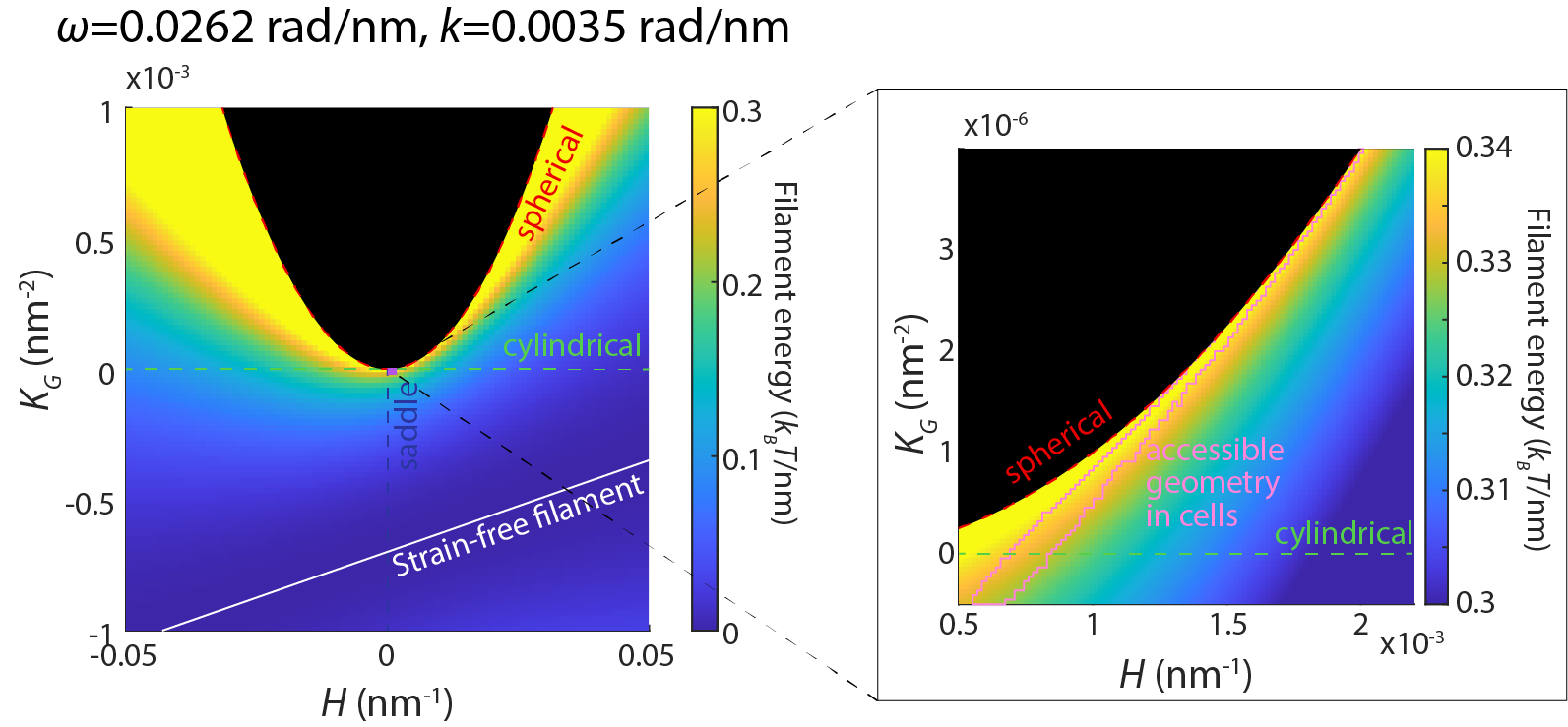

**Figure S3: Fitting of MreB experimental data to model.** Due to the geometric defined by the relatively rigid cell body, MreB filaments deform from a strain-free conformation to bind to the membrane. Right: zoomed-in version highlighting the region of geometries accessible within cells. Note the difference in color bar scales between the two panels. Black regions: geometries that are impossible.

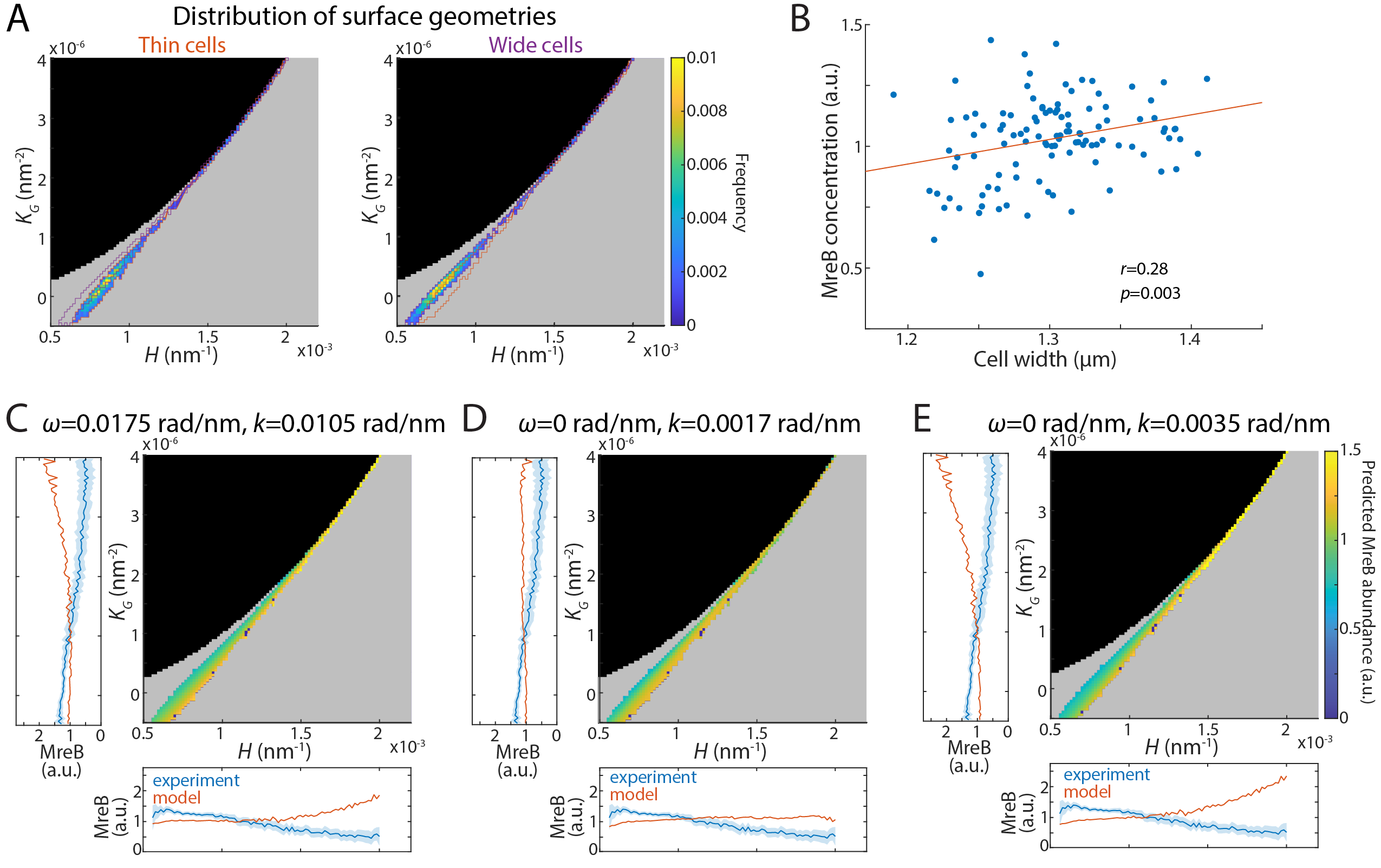

**Figure S4: The model fails to recapitulate experimental results at large *k* and small *ω*.**

A) The distribution of cell contour geometries along thin (left, cell widths in bottom 30^th^ percentile) and wide (right, cell widths in top 30^th^ percentile) cells. Thin and wide cells occupy distinct spaces in terms of mean and Gaussian curvatures. Red (purple) outlines denote the distribution of geometries in the thin (wide) cell populations, respectively.
B) MreB concentration is correlated with cell width. *n*=193 cells. Pearon’s *r*, *p*-value was calculated using a two-tailed Student’s *t*-test.
C) Model fitting of experimental data in Fig. 4A. For this set of *ω* and *k* values, the model predicts higher MreB abundances at higher mean (lower plot) or Gaussian (left plot) curvatures. Blue shaded regions denote 95^th^ confidence intervals estimated by bootstrapping.

D,E) When a filament exhibits no intrinsic twist (*ω* = 0), the model generally predicts higher MreB abundances at higher mean (lower plot) or Gaussian (left plot) curvatures, inconsistent with experimental data. Blue shaded regions denote 95^th^ confidence intervals estimated by bootstrapping.

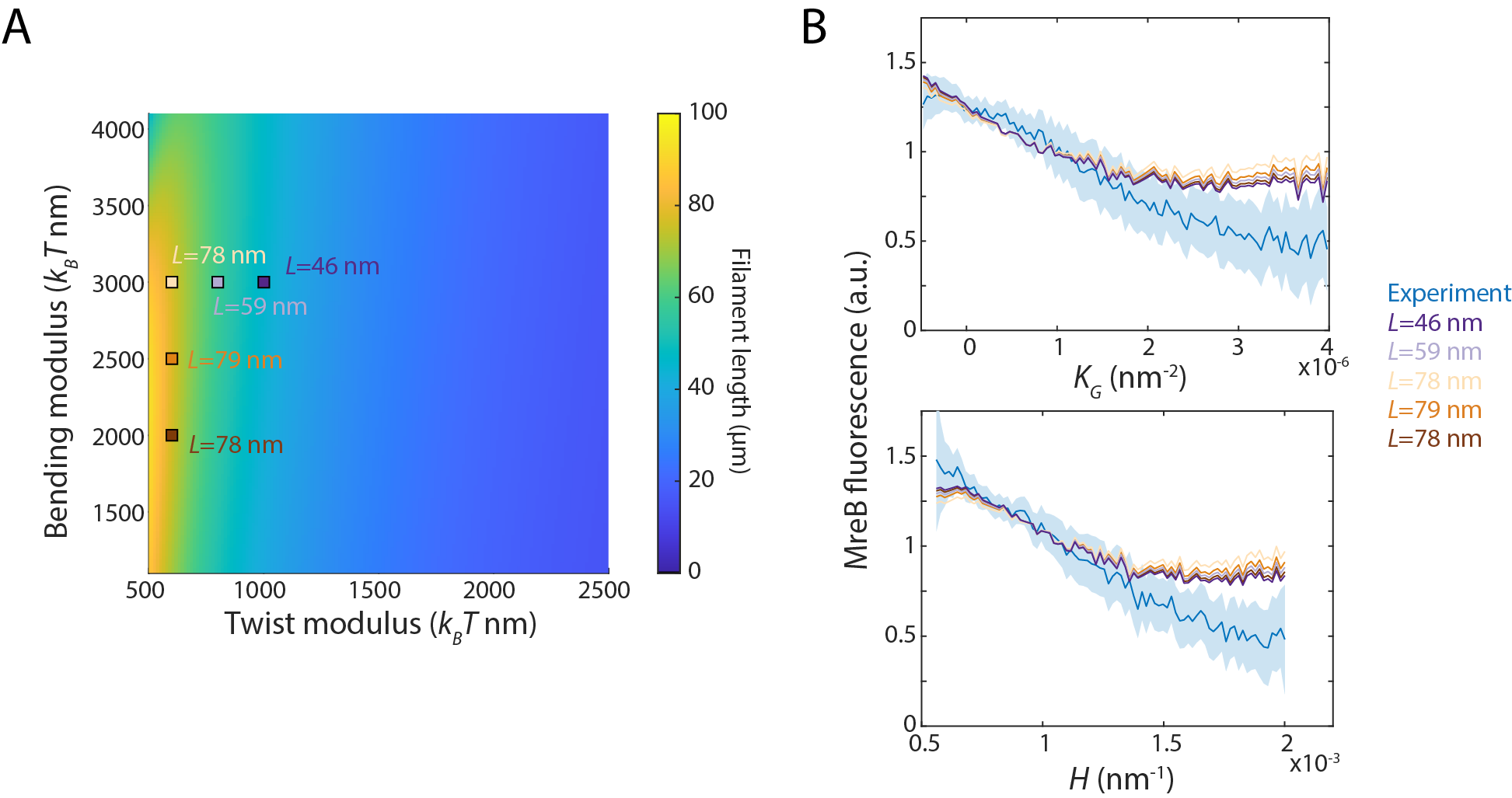

**Figure S5: Model fitting is robust to changes in twist and bending moduli.**

A) Predicted MreB filament length across twist and bending moduli. Five parameter sets spanning estimates from MD simulations are highlighted by squares.
B) Fits corresponding to the 5 parameter sets highlighted in (A). The energy landscape remained similar across estimates of twist and bending moduli, and the model predicted curvature-dependent MreB localization consistent with experimental data.
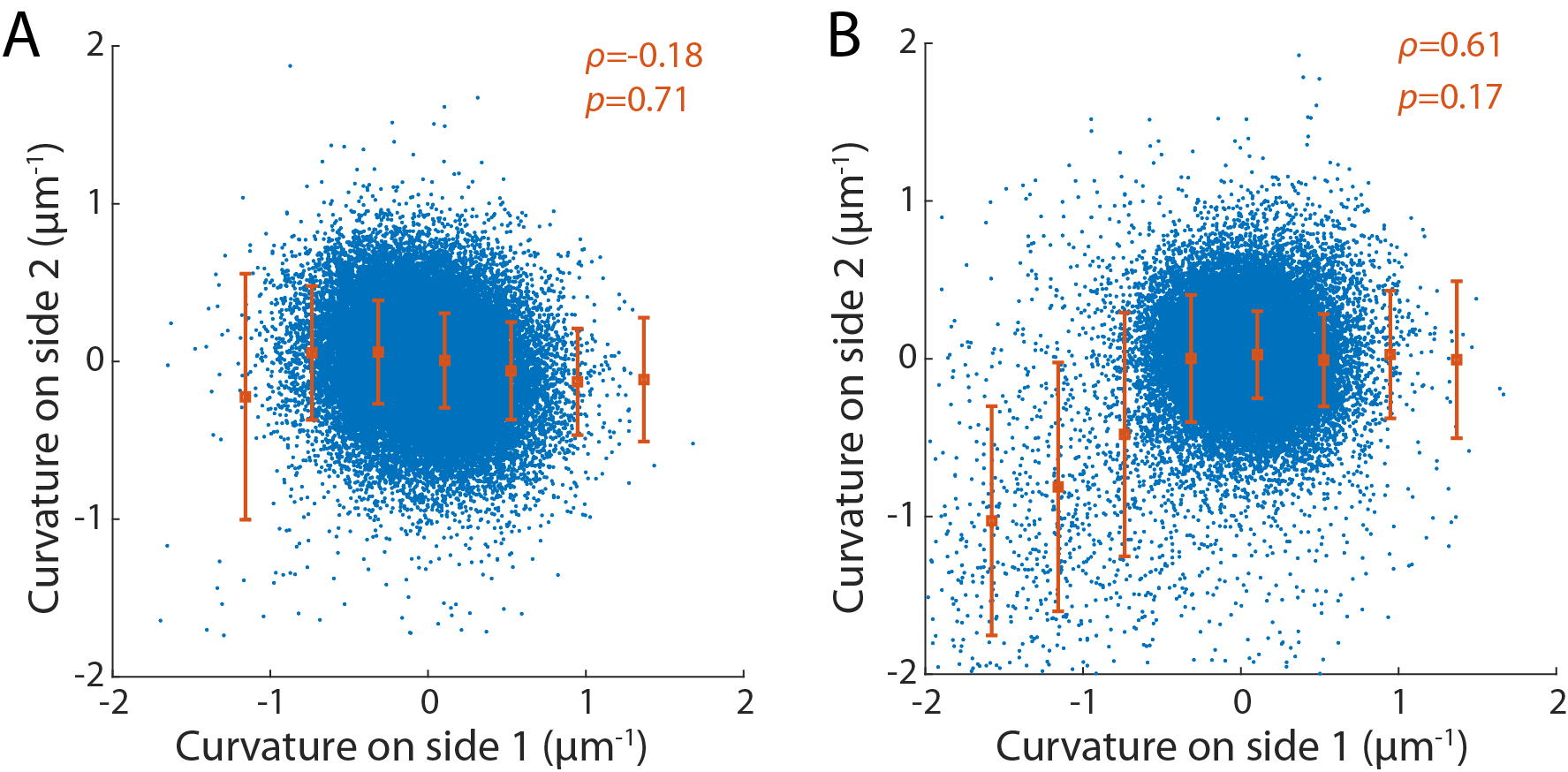

**Figure S6: Contour curvature is uncorrelated with the opposing point relative to the cell midline.**

In-plane contour curvatures along the cell body for cells with (A) or without (B) treatment with the division inhibitor cephalexin. For paired points on opposite sides of the midline, the curvature on one side was uncorrelated with the curvature on the other side. Blue dots: raw data; red circles: mean of binned curvature values, error bars represent 1 S.D. *ρ*: Spearman’s correlation coefficient of binned data. *p*-values were calculated from two-tailed Student’s *t* tests.
